# Termite communities along a disturbance gradient in a West African savanna

**DOI:** 10.1101/167692

**Authors:** Janine Schyra, Judith Korb

**Author notes:** corresponding author. Uni-Osnabrueck.de.

## Abstract

Termites are important ecosystem engineers, crucial for the maintenance of tropical biodiversity and ecosystem functioning. But they are also pests which cause billions of dollars in damage annually to humans. Currently, our understanding of the mechanisms influencing species occurrences is limited and we do not know what distinguishes pest from non-pest species. We analyzed how anthropogenic disturbance (agriculture) affects species occurrences. We tested the hypothesis that strong disturbance functions as a habitat filter and selects for a subset of species which are major pests of crop. Using a cross-sectional approach, we studied termite community composition along a disturbance gradient from fields to 12-year-old fallows in a West African savanna. We reliably identified 19 species using genetic markers with a mean of about 10 species - many of them from the same feeding type - co-occurring locally. Supporting our hypothesis, disturbance was associated with environmental filtering of termites from the regional species pool, maybe via its effect on vegetation type. The most heavily disturbed sites were characterized by a subset of termite species which are well-known pests of crop. This is in line with the idea that strong anthropogenic disturbance selects for termite pest species.

## Introduction

Termites are major ecosystem engineers with crucial roles in decomposition, soil fertility, hydrology, and species diversity (Pringle et al. 2010; Evans et al. 2011). Concomitantly, a few species are also major pests (Rouland-Lefèvre 2011). Despite their importance, we hardly understand what determines the occurrence of different termite species and what distinguishes pest from non-pest species. Niche overlap between different species seems to be substantial as termites are detritivores and only four major feeding types are distinguished (Donovan et al. 2001): dead wood feeders (group I); dead wood, micro-epiphytes, leaf litter and grass feeders (group II); humus feeders (group III) and true soil feeders (group IV) (reviewed in (Davies et al. 2003; Eggleton 2011). In African savannas up to 20 higher termite species (Termitidae) of feeding group II co-exist (e.g., Dosso et al. 2010, 2013; Hausberger et al. 2011; Hausberger and Korb 2015, 2016). These group II species can be subdivided into two feeding type specialists, grass-feeding *Trinervitermes* (group II_g_) and fungus-growing Macrotermitinae (group II_f_). The latter cultivate an obligate symbiotic fungus within their colonies, which they provision with a broad range of dead plant material (Nobre et al. 2011). The similarity of termites’ food niches implies that competitive interactions are important in shaping local savanna communities (Basu 2011; Korb and Linsenmair 2001a). However, recent analyses suggest that random processes play an important role in community assembly in an undisturbed West African savanna, with a structuring effect by one large mound building species, *Macrotermes bellicosus* (Hausberger and Korb 2015). Additionally, first evidence implies that assembly processes change to more environmental filtering with disturbance (Hausberger and Korb 2016). This suggests that disturbance can not only lead to a decline in species richness, but also to a change of the processes that structure species communities.

In the current study, we investigated termite community composition of different-aged fallows (measured as the time since they were last cultivated) in a West African savanna region in Togo. By doing this, we aimed at analysing how communities assemble over time from a strong anthropogenically disturbed habitat to a more natural setting. We expect that termite communities of younger-aged fallows are more strongly structured by environmental filters. To test this, we first identified all species occurring in the different communities using morphological and genetic markers. A genetic approach is necessary to unambiguously identify all termite samples. To reveal assembly processes, we then applied phylogenetic community analyses that explicitly test real, studied communities against communities that are drawn at random from the regional species pool. Finally, we investigated whether species from young-aged fallows correspond to pest species.

## Materials and Methods

### Termite sampling

Termites were systematically collected when they were most active, that is, during the beginning of the rainy season, near the Oti-Keran National Park in northern Togo (West Africa; 10°17’ to 10°08’ N; 0°28’ to 0°51’ E, Fig. 1). This region is a typical West African savanna lying in the center of the West Sudanian biome (mean annual precipitation: 1100 mm; Worldclim database). Termites were collected in 2012 from seven fallows of age 0, 2, 4, 6, 8, 10 and 12 years. In 2014 we added six new fallows of age 0, 0, 1, 2, 10 and 10 years. Our sampling regime was constrained by the availability of fallows with known age.

**Fig. 1.**
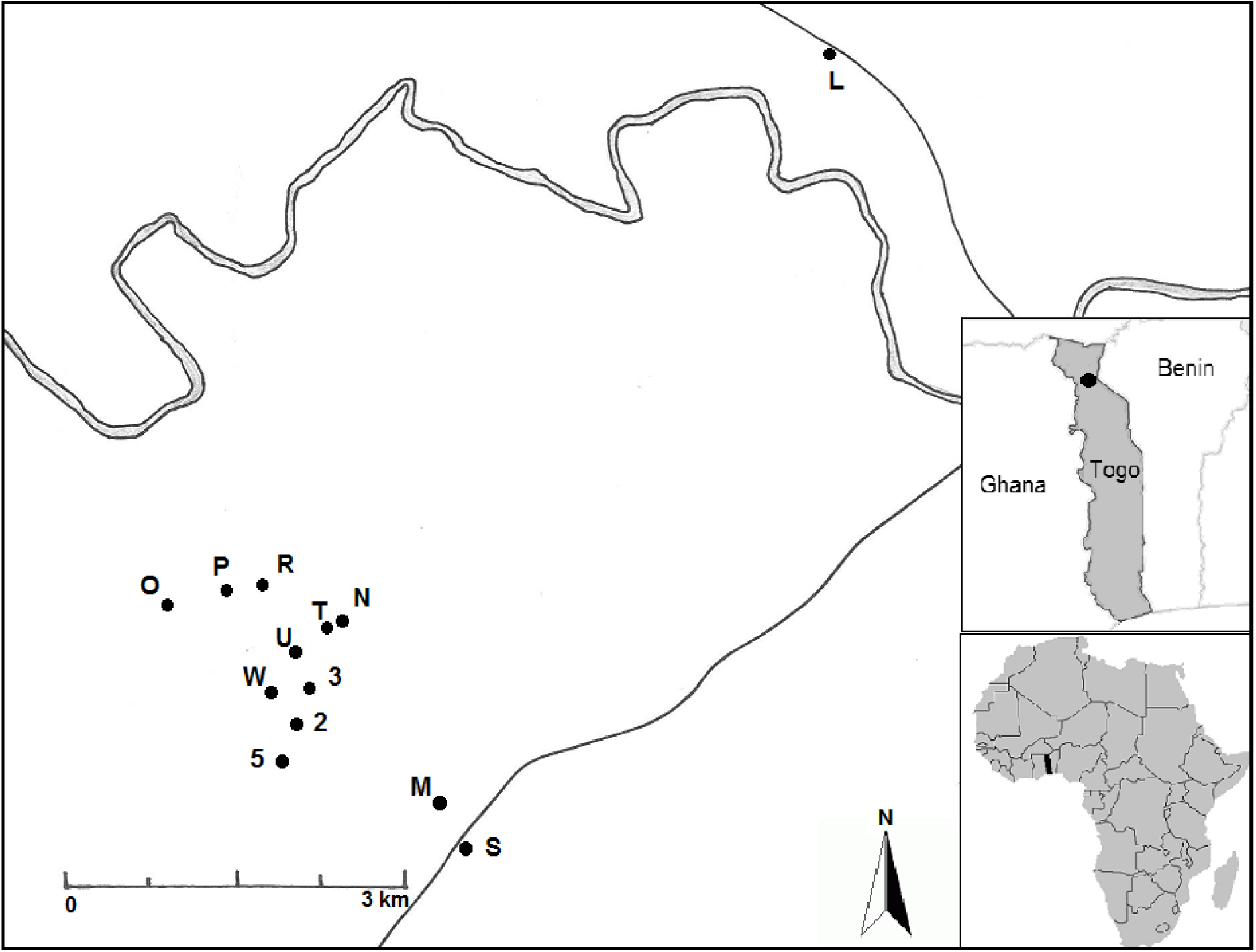
Location of the the Oti-Keran National Park in northern Togo and distribution of the 13 sampled fallows in the study area. Single lines are small roads and the double line in grey is a river

Sampling was done using a standardized belt transect protocol first developed for sampling termites in forests (Jones and Eggleton 2000) and then adapted to savannas (Hausberger et al. 2011). In short, the protocol consists of a thorough search of dead plant material on the ground, on and in trees and mounds as well as soil sampling to assess termite diversity (Jones and Eggleton 2000). Plot size was one hectare with three transects each measuring 2 m x 50 m, divided into ten 2 m x 5 m sections, arbitrarily located within one plot. Each transect section was searched systematically for termites for 15 minutes by a trained person. Additionally, we sampled eight soil scrapes per transect section measuring 15 cm x 15 cm x 10 cm. All encountered termites were stored in 99% pure ethanol for subsequent molecular analyses.

As in the former studies (Hausberger et al. 2011; Jones and Eggleton 2000; Korb and Linsenmair 2001b), we chose a plot size of one hectare because the foraging ranges of termite colonies is within 100 m (Korb and Linsenmair 2001b). Hence one hectare represents the local scale where interactions between colonies occur, i.e., it reflects the Darwin-Hutchinson-Zone, which is most relevant to study assembly of local communities (Vamosi et al. 2009). We specifically selected plots with and without active *M. bellicosus* mounds as it is the main mound builder and an important ecosystem engineer which may influence termite communities.

All samples were identified to the species level: Samples containing soldiers were first identified using the keys by Webb (1961) and Sands (1965a), and then sequenced to obtain an unambiguous species identity (see below). Samples with workers only, were genetically analysed, as morphological identification was impossible (see below). To assign the feeding group to each sample, we followed the anatomical criteria outlined by Donovan et al. (2001). Whenever we found/encountered termites during the search within a transect section, we collected a few specimens in a vial (5-10 individuals). Then we continued searching within the section and when we encountered termites again they were placed in a separate vial. The number of all resulting vials for a study plot (i.e., the sum over all transect sections for all three replicate transects within a plot) was used as encounter rate. This is used as a surrogate of species abundance (Davies 2002). Naturally, the presence / absence of each species within each plot arose from these data as well.

During sampling, we recorded data for the environmental variable ‘vegetation type’ by classifying plots according to their vegetation type and density: field (recognizable cultivation and crop plants), open savanna (mainly grass land, few bushes and trees), and medium-dense savanna (many bushes and trees). The savanna was a typical West African Sudanian savanna. The main shrub and tree species were *Afzelia africana, Crossopteryx febrifuga, Detarium microcarpum, Piliostigma thonningii, Vitellaria paradoxa, Combretum* spp., *Terminalia* spp. and *Gardenia* spp. All study plots were located away (at least 1 km) from rivers or lakes. The topography of the studied plots was flat.

### Genetic identification and phylogenetic analyses

To allow unambiguous species identification, we isolated DNA and sequenced fragments of three genes as described elsewhere (Hausberger et al. 2011) (additional data are given in Online Resource 1: Table S1): *cytochrome oxidase subunit I* (*COI*; total length 680bp), *cytochrome oxidase subunit II* (*COII*; total length 740 bp), and *12S* (total length 350 bp). These sequences were used to re-construct phylogenetic trees using three approaches (Bayesian method, maximum-parsimony analysis, and maximum-likelihood analysis) to delimitate and identify species (for more details see Supplementary). As in former termite studies (Legendre et al. 2008; Hausberger et al. 2011), *COII* was most useful for ‘barcoding’ (i.e., assigning species to samples) because it amplified well and gave appropriate resolution for species identification. All samples were identified. Species names correspond to those given in Hausberger et al. (2011) and Hausberger and Korb (2015, 2016) for Benin. *Amitermes* sp1 from the Benin studies is actually *Amitermes evuncifer*. Hence we used the proper species name in our study. To obtain corresponding species identities, we constructed a phylogeny comprising all species occurring in Togo and Benin. Samples forming a species cluster were named identically. In total we sequenced 899 samples in the current study.

### Phylogenetic community structure analyses

We analysed the local community structure with PHYLOCOM 4.2 (Webb et al. 2008). As input tree for the phylogenetic community structure analyses we used the *COII* gene tree, which was pruned prior to analysis so as to have only species of the regional species pool included and only one representative per species in the tree. This representative was the sequence with the highest quality values for each base (maximum value of 61, multiplied by ten) as defined in Chromas 2.4.4 (1998-2016, Technilysium Pty Ltd).

We calculated the net relatedness index (NRI) that measures whether locally co-occurring species are phylogenetically more / less closely related than expected by chance. It uses phylogenetic branch length to measure the distance between each sample to every other terminal sample in the phylogenetic tree, and hence the degree of overall clustering (Webb et al. 2008). The NRI is the difference between the mean phylogenetic distance (MPD) of the tested local community and the MPD of the total community (regional) divided by the standard deviation of the latter. High positive values indicate clustering; low negative values overdispersion (Webb et al. 2002). We tested whether our data significantly deviated from 999 random communities derived from null models using the independent swap algorithm on presence / absence data (Gotelli and Entsminger 2003; Hardy 2008). The swap algorithm creates swapped versions of the sample / species matrix and constrains row (species) and column (species’ presence or absence) totals to match the original matrix. The regional species pool consisted of all species from all studied localities. As suggested by Webb et al. (2008), we used two-tailed significance tests based on the ranks that describe how often the values for the observed community were lower or higher than the random communities. With 999 randomisations, ranks equal or higher than 975 or equal and lower than 25 are significant at *P* ≤ 0.05 (Bryant et al. 2008).

### Similarity between fallows

We quantified the compositional similarity (ß-diversity) between all localities using the Bray-Curtis sample similarity index (Magurran 1988), which was calculated in EstimateS version 8.2.0 (Colwell 2013). It ranges from 0 to 1, with low values indicating low similarity and high values the reverse. The Bray-Curtis index is quantitative; the abundance of species is taken into account when calculating the shared species statistics.

### Other statistical analyses

All inferential statistics were done with the statistical package IBM SPSS 16. All tests were two-tailed. Data were tested for assumptions of parametric testing and analyses were done accordingly. For all data, qualitatively the same results (i.e. effects were significant or non-significant) were obtained when testing parametrically or non-parametrically.

## Results

### Diversity

We identified a total of 19 termite species (regional species pool), all Termitidae (Table 1). All conducted phylogenetic analyses yielded similar topologies (Fig. 2; Online Resource 1: Fig. S1, S2). As is typical for African savannas, fungus-growing Macrotermitinae dominated with eight species (*Microtermes* sp.1, *Microtermes* sp.2, *Microtermes* sp.3, *Microtermes* sp.4, *Macrotermes bellicosus, Macrotermes subhyalinus, Ancistrotermes* sp.1, *Odontotermes* sp.1), of which all co-existed locally. The next most-species rich group were Nasutitermitinae with the grass-feeders *Trinervitermes occidentalis, Trinervitermes geminatus, Trinervitermes oeconomus, Trinervitermes togoensis* and *Fulleritermes tenebricus*. Despite occupying the same feeding niche, all four *Trinervitermes* species co-occurred locally. Further, we sampled two representatives of the soil-feeders Apicotermitinae (*Astalotermes* sp., *Adaiphrotermes* sp.1) and four species of the Termitinae (*Microcerotermes* sp.1, *Amitermes evuncifer* (both wood-litter feeders), *Procubitermes* sp.1 and *Pericapritermes* sp.1 (soil and soil-wood feeders)). Surprisingly, species richness did not increase with fallow age (Spearman-rank correlation: N = 13, P = 0.324). Out of a total of 19 species that we found in the fallows, from seven to 13 species co-occurred locally. Species names are in accordance with the identified species from the studies in Benin (Hausberger et al. 2011; Hausberger and Korb 2015, 2016).

**Fig. 2.**
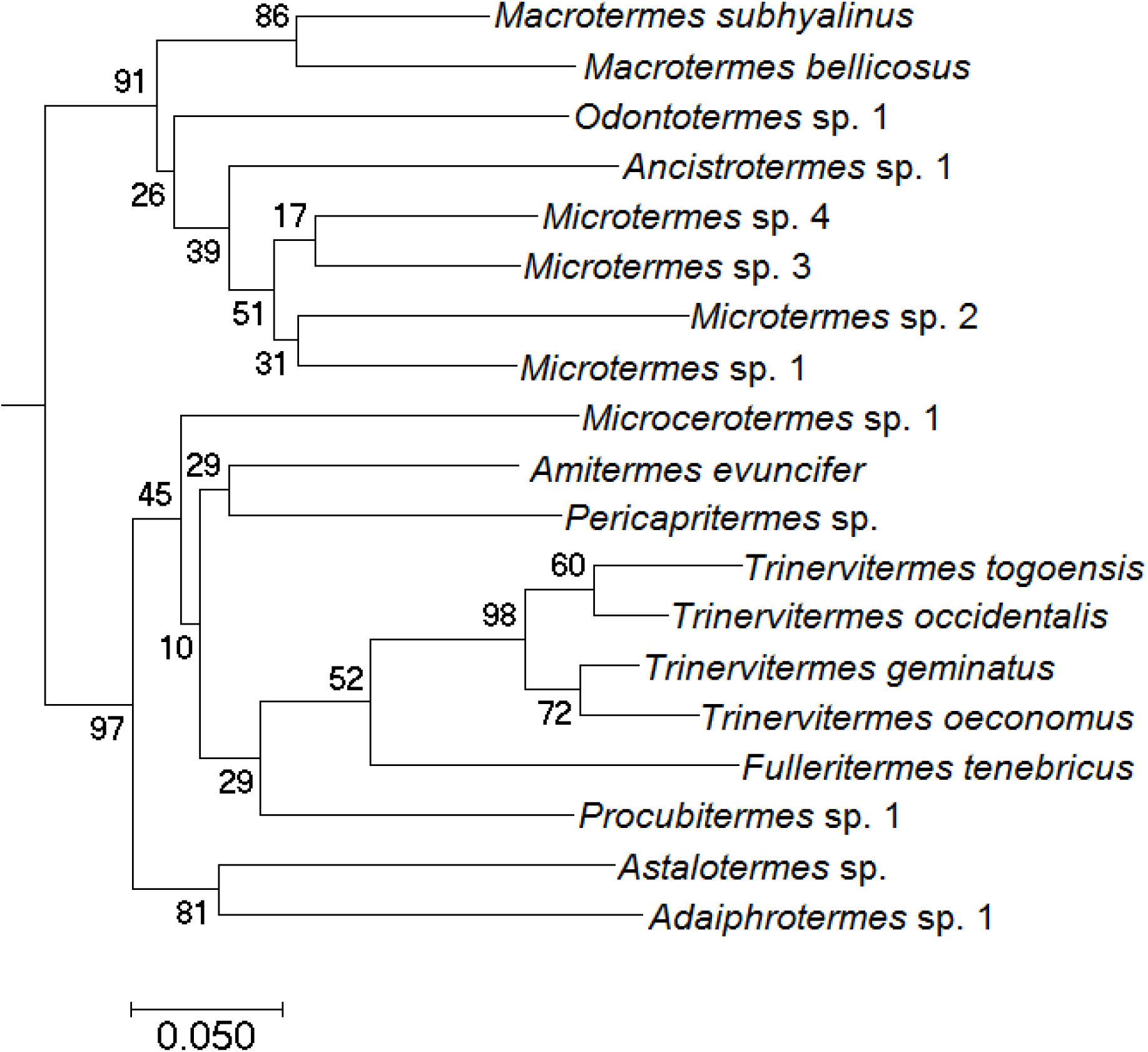
Input Bayesian phylogeny for Phylocom based on the gene cytochrome oxidase II using MrBayes v3.1.2. Analysis was done with 10^7^ generations, number of chains=4, sample frequency=1000 and a finalizing burn-in of 2500. Node numbers are the posterior probabilities calculated to assess branch support

**Table 1.**
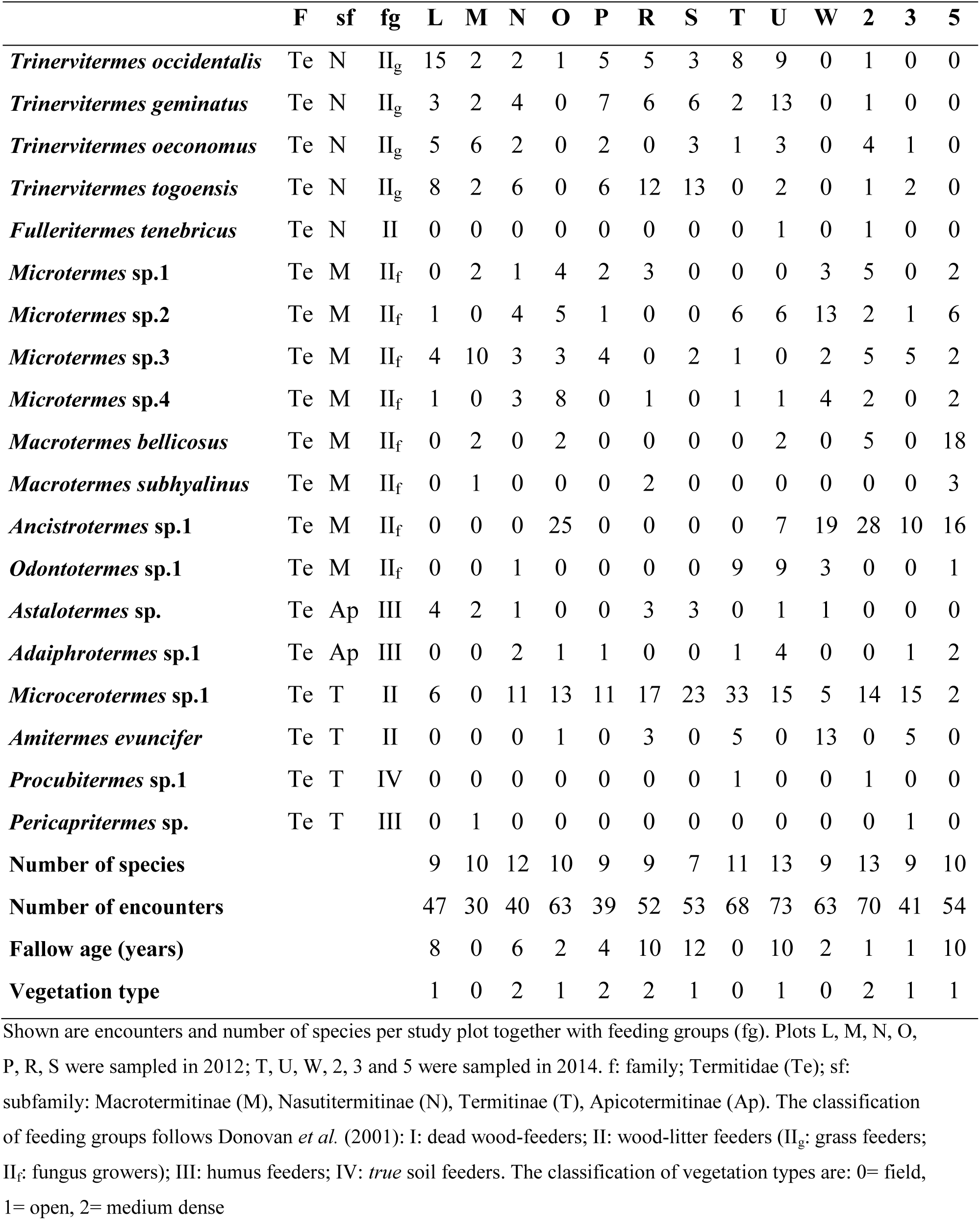
Abundances and encounters of the 19 species of the regional species pool in all study plots including data on fallow age and vegetation type

### Phylogenetic community structure

The NRI values, measuring the phylogenetic community composition, ranged from −0.72 to 4.21. Three plots showed significant signals of environmental clustering (Plot S: NRI: 2.83; Plot W: NRI: 2.80; Plot 5: NRI: 4.41; all P < 0.05). NRI values did not correlate with fallow age (Spearman-rank correlation: N = 13, P= 0.131) nor with species richness (Spearman-rank correlation: N = 13, P = 0.890). However, there was an indication that vegetation type affects phylogenetic community structure (ANOVA: F = 3.21, P = 0.084, Fig. 3). Communities tended to be more phylogenetically clustered in fields and especially open savannas (Turkey HSD *post-hoc* test: *field/open*: P = 0.802; *field/medium dense*: P = 0.316; *open/medium dense*: P = 0.072. Similarly, *M. bellicosus* may have an effect on phylogenetic structuring: as a tendency, when *M. bellicosus* was present NRIs were higher (i.e. more phylogenetically clustered communities) than when it was absent (Mann-Whitney-U test, Z = −1.76, N = 13, P = 0.079; Fig. 4).

**Fig. 3.**
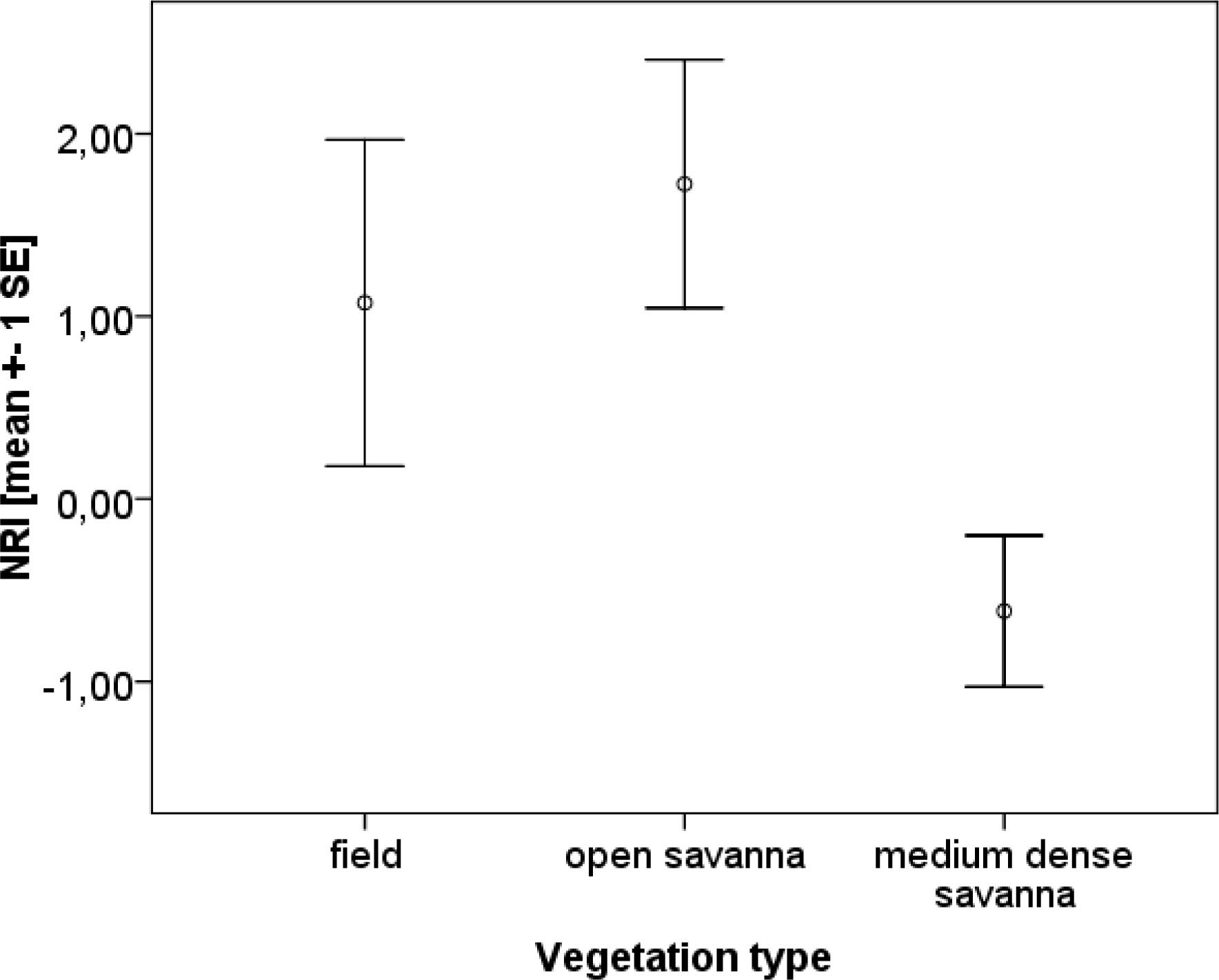
NRI (Net Relatedness Index) and vegetation type. There was an indication that vegetation patterns affect termite community composition, as communities were more clustered (more closely related) in open savannas

**Fig. 4.**
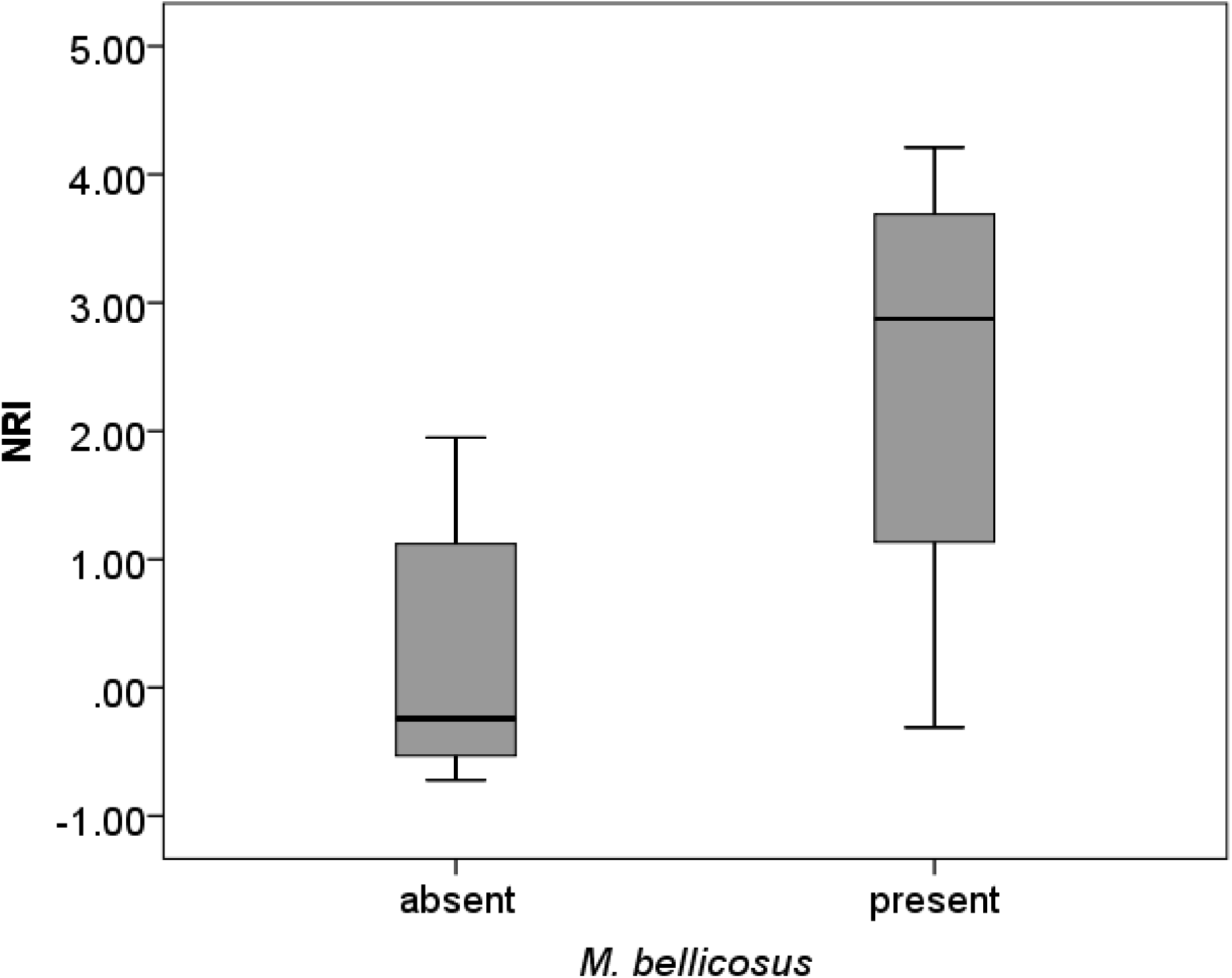
NRI and presence of *M. bellicosus*. There may be an effect of *M. bellicosus* on phylogenetic community structure. NRIs were higher (i.e. more phylogenetically clustered communities) when *M. bellicosus* was present than when it was absent (P=0.079)

### Similarity between fallows

The compositional similarity between sites varied, with the Bray-Curtis index ranging from 0.075 to 0.785. Mean species richness per site was 10.1 (± SD 1.75) species and mean number of shared species between sites was 6.1 (± SD 1.64) species. When comparing sites of different vegetation types to each other, the Bray-Curtis index revealed that there is a significant difference in species composition between vegetation types (ANOVA: F = 4.329, P = 0.002, Fig. 5). Fields (f/f) were significantly less similar among each other in species composition than open compared to medium dense savanna sites (o/m) are, or medium dense savanna sites (m/m) are among each other (Tukey-HSD *post-hoc* test: f/f vs o/m: P = 0.022; f/f vs m/m: P = 0.016). Open savanna sites and medium dense savanna sites had a higher species similarity among and between each other. The other compared vegetation types (f/o, f/m, o/o) lay between these extremes. Overall, there was a pattern that compositional species similarity rises, the less disturbed the sites are.

**Fig. 5.**
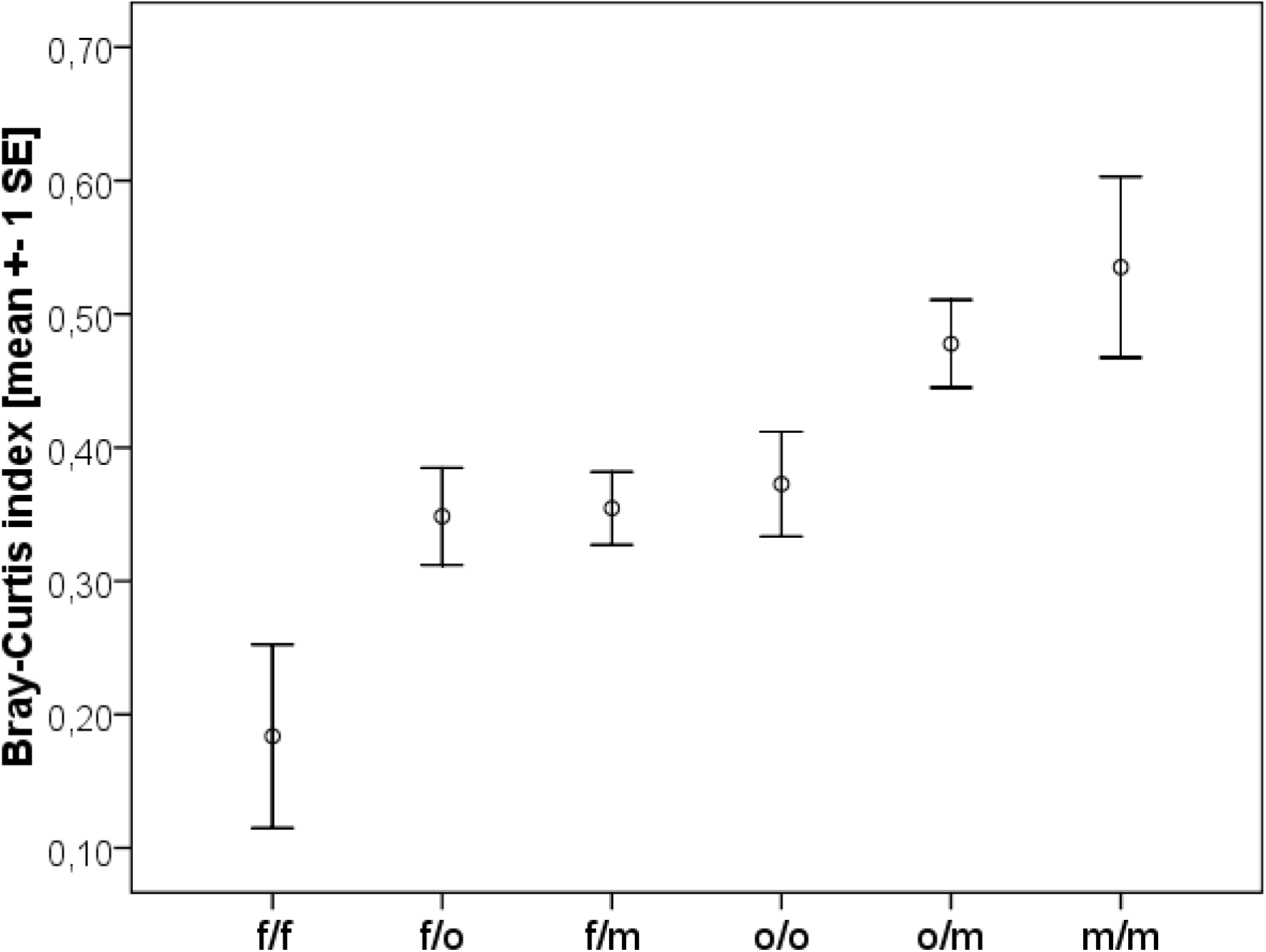
Compositional species similarity between plots of different vegetation type measured with the Bray-Curtis sample similarity index. It revealed a significant difference in species similarity between sites of different vegetation types

### Pest species

We could identify several termite species in our study sites that are known pest species (Wood et al. 1980; Rouland-Lefèvre 2011; Cowie et al. 1989; Collins 1984): *M. subhyalinus, Odontotermes* sp., *Microtermes* sp., *Pericapritermes sp., Amitermes evuncifer* and *Ancistrotermes* sp.. We found all these species throughout all fallows. Yet, especially *Microtermes* species, *Amitermes evuncifer* and *Ancistrotermes* sp. were quite common in young fallows and *Pericapritermes* sp. was only sampled in young fallows (0-2 years; vegetation type ‘field’, Table 1). For *Odontotermes* sp. and *Macrotermes subhyalinus* we did not find a specific pattern. These species occurred in young and older fallows alike, but these species were rare in general.

### Discussion

Our results support the hypothesis that strong anthropogenic disturbance is associated with environmental filtering of termite species in savannas. We found no evidence for interspecific competition playing a major role in structuring these termite communities.

Similar to the study on termite communities of the West African savanna in Benin (Hausberger and Korb 2016), we found a total of 19 termite species in this study in Togo, with 17 (89.47%) species shared between both regions. Despite their similar food requirements, on average 10 species co-occurred locally, all belonging to the higher termites and often sharing the same feeding and sub-feeding type. Based on the phylogenetic community analysis, we found no evidence that interspecific competition is playing a major role in structuring these communities. On the contrary, especially in strongly disturbed sites, environmental filtering seems to be important (Fig. 3, 5). This confirms what was found for Benin and implies that disturbance is associated with environmental filtering in West African savanna regions.

In contrast to Benin, we did not detect a decline of species richness with disturbance (Hausberger and Korb 2016). Rather a higher degree of disturbance in younger fallows seems to favour a certain set of species. This difference between both studies may be due to the drier climate in Benin and /or because in Benin samples were taken next to villages with ongoing land-use. Many of the species from fields are known as crop pests in West Africa, especially *Amitermes evuncifer, Ancistrotermes* sp., *Microtermes* sp. (Table 1) (Wood et al. 1980; Rouland-Lefèvre 2011; Cowie et al. 1989; Collins 1984). Whether the occurring termite species are pests because they are more resilient against disturbance or whether selection as pests made them more resilient is difficult to test. As some of the sampled pest species also occurred in some older fallows, in particular *Odontotermes* sp. and *Macrotermes subhyalinus*, this suggests that they are generalists that can cope better with human disturbance. By contrast, grass-feeding *Trinervitermes* species (*T. togoensis, T. geminatus, T. oeconomus* and *T. occidentalis*) were mainly found in older fallows (Table 1), implying that they are less resilient against disturbance. As grass is also commonly available in young fallows, it is unlikely that limited food availability can account for their absence. Nevertheless, fields seem to be very heterogeneous among each other concerning species composition as indicated by the low Bray Curtis similarity (Fig. 5). This could be due to the kind of crop being cultivated or other biotic and abiotic factors.

As in Benin (Hausberger and Korb 2015), several closely related fungus growers were associated with the occurrence of *M. bellicosus* (including *Microtermes* spp., *Ancistrotermes* sp.1 and *Macrotermes subhyalinus*), reflected in phylogenetic clustering (Fig. 4). *Macrotermes* mounds can provide micro-habitats for other fungus growers as well as facilitating their occurrence by concentrating nutrients and clay through their nest building and foraging activities (Joseph et al. 2013), thereby explaining the increased phylogenetic clustering in sites with *M. bellicosus* mounds.

### Comparison with other community studies in West Africa

There are few studies on West African termite communities and besides the above mentioned recent studies in Benin, none used a molecular approach necessary for unambiguous species identification of West African termites, which makes direct comparisons difficult.

Dosso et al. (2013) studied termite communities in a more wooded region near Lamto in Ivory Coast in land-use systems ranging from a semi-deciduous forest, over plantations, to a crop field and a 4-year old fallow. As is typical for forests (Eggleton et al. 2002), species richness declined - and especially soil feeders disappeared - with disturbance and a transition from forest to a more open habitat. The crop field and the 4-year old fallow, which are most comparable to our study sites, harboured 11 and 7 morpho-species, respectively. Several of these species are typical forest species and hence absent in our study (e.g. *Nasutitermes, Basidentitermes*). Only a single *Microtermes* species was found in Lamto, compared to four in this study and in Benin (Hausberger et al. 2011). One species might be an underestimation as *Microtermes* species are difficult to identify without genetic means. Strikingly, only one *Trinervitermes* species was found in Lamto and this in the 4-year old fallow. This supports our conclusion that *Trinervitermes* spp. are susceptible to disturbance. Another study near Lamto tested the influence of annual fires on termite diversity (Dosso et al. 2010). Here, the occurrence of *Trinervitermes* spp. in burnt areas decreased, further supporting the hypothesis that they are less resilient species.

In contrast to our study, some studies implicated evidence for interspecific competition in structuring termite communities (Su and Scheffrahn 1988; Leponce et al. 1996; Korb and Linsenmair 2001a; Bourguignon et al. 2009, 2011; Li et al. 2015; Basu 2011). Reasons for these diverse conclusions include differences between study sites, disturbance regimes, and lack of testing against the null hypothesis of random community assembly. Additionally, most studies focused on a few species only and did not study whole termite communities, therefore addressing a different scale. More studies, spanning more regions, are necessary to derive general conclusions. Such studies should cover complete communities of genetically identified species where species co/occurrences are tested against random assemblages. Genetic identification is helpful as otherwise especially the most closely related species may be misidentified, which can lead to blurring signals of environmental filtering or interspecific competition.

## Acknowledgements

We thank the Université de Lomé in Togo, especially Jean Norbert Gbenyedji, Boris Dodji Kasseney, Banibea Sanbena Bassan, and the local villagers on-site, for substantial help during field work and logistic support. The project was funded by the Deutsche Forschungsgemeinschaft (DFG) (Project KO1895/12-1).

